# Chemical composition and insecticidal activity of *Aeollanthus pubescens* Benth leaf essential oil on the malaria vector *Anopheles gambiae* (Diptera: Culicidae)

**DOI:** 10.1101/2020.11.25.397588

**Authors:** Roméo Barnabé Bohounton, Luc Salako Djogbénou, Oswald Yédjinnavênan Djihinto, Oronce Sedjro-Ludolphe Dedome, Pierre Marie Sovegnon, Bruno Barea, Aristide Adomou, Pierre Villeneuve, Fidèle Paul Tchobo

**Author notes:** **Corresponding author Mail:** (LSD).

## Abstract

The use of synthetic insecticides is responsible for many cases of resistance in insects. Therefore, the use of natural molecules of ecological interest with insecticidal properties turns out to be an alternative approach to the use of synthetic insecticides. This study aims at investigating the larvicidal, adulticidal activity and the composition of the essential oil of *Aeollanthus pubescens* Benth on the major malaria vector *Anopheles gambiae*.

The leaves of *Aeollanthus pubescens* were collected in the South of the Republic of Benin. Three reference strains of *Anopheles gambiae s*.*s*. such as Kisumu, Kiskdr and Acerkis were used. The chemical composition of the essential oil was analysed by gas chromatography coupled to mass spectrometry. Larvae were exposed to the essential oil extract for 24 h. Adult mosquitoes were exposed to the fragment nets coated with the essential oil for 3 min. Larval mortality and adult survivorship were monitored.

Fourteen components were identified representing 98.31% of the total of oil. The major components were carvacrol (51.06 %), thymyle acetate (14.01 %) and γ-terpinene (10.60 %). The essential oil has remarkable larvicidal properties with LC_50_ of 29.26, 22.65, and 28.37 ppm respectively on Kisumu, Acerkis and Kiskdr strains. With the fragment net treated at 165 µg/cm^2^, the KDT_50_ of both Acerkis (1.71 s, *p* < 0.001) and Kiskdr (2.67 s, *p* < 0.001) individuals were significantly lower than that of Kisumu (3.77 s). The lifespan of the three mosquito strains decreased respectively to one day for Kisumu (*p* < 0.001), two days for Acerkis (*p* < 0.001) and three days for Kiskdr (*p* < 0.001) compared to their control.

Our findings show that the *Aeollanthus pubescens* essential oil is an efficient larvicide and adulticide against malaria vector *Anopheles gambiae*. This bioinsecticidal activity is a promising discovery for the control of the resistant malaria-transmitting vectors.

## Introduction

Vector-borne diseases remain the major causes of death in many tropical countries. The most important vector-borne diseases are malaria, lymphatic filariasis, dengue fever and yellow fever; the pathogens responsible for these diseases are transmitted by the arthropods vectors including mosquitoes (Diptera: Culicidae) which are the major agents [1,2]. Among the mosquito-borne infectious diseases, malaria is the most dreadful and the major public health concern in terms of the number of incidence, prevalence, morbidity and mortality in low-income countries of Africa, Asia, Latin America and beyond [3,4]. Despite the national malaria control programmes efforts, nearly 85% of malaria deaths occurred in 21 sub-Saharan African countries including the Republic of Benin [4]. Malaria is transmitted by the bites of parasite-infected *Anopheles* female mosquitoes that carry out the infection to humans [5].

So far, most malaria control programs have mainly relied on Artemisinin-based Combination Therapies (ACTs) for the treatment of diagnosed patients and the use of chemical compounds through Insecticides Treated Nets (ITNs) and Indoor Residual Spraying (IRS) for the prevention of human-vector contacts [6]. Nowadays, 14 insecticides belonging to four major classes of synthetic chemical insecticides are recommended by WHO Pesticide Evaluation Scheme (WHOPES) for IRS [7] and four insecticides, all from the pyrethroids group, are recommended for ITNs [8]. However, the overuse of these synthetic chemical insecticides has led to the emergence of resistant malaria vectors; and the frequency of insecticide resistance is widespread especially in African regions [9,10]. Also, these synthetic insecticides have been recognized to have adverse effects on non-targeted species and affect animal and plant biodiversity [11–13]. The indiscriminate use of these chemicals has also been shown to have severe effects on the environment and impacts on human health [14].

These facts combined with multidrug-resistance in malaria parasite [15] and the absence of an effective vaccine led scientists to focus on searching for environmentally-friendly vector control alternatives with the aim of decreasing the selection pressure for insecticide resistance [16]. This kind of eco-friendly vector control alternatives could be achieved especially with insecticide from botanical sources. Indeed, they are potentially safer for human and the environment and have a minimal residual effect. They are more target-specific; less toxic to vertebrates, and more sustainable than their synthetic conterparts [17,18].

Beninese traditional medicine and pharmacopoeia medications are richly bio-diversified, which could be a great source for natural insecticides for malaria control [19]. Therefore, it appeared of interest to learn more about Beninese flora for its insecticidal activities. However, to our knowledge, only few studies have been conducted regarding the mosquitocidal activity of the extracts from Beninese plants species [20–22].

*Aeollanthus pubescens* (*A. pubescens*) Benth is an annual herbaceous plant distributed in many West African countries as well as in Benin [23]. It is commonly used by local populations as food and also for it medicinal properties [24,25]. The current study aims at determining the insecticidal potential of *A. pubescens* Benth leaf essential oil against Afrotropical malaria vector *Anopheles gambiae* under laboratory conditions by investigating safer alternatives for the management of malaria vector resistance compared to existing synthetic insecticides.

## Material and methods

### Plant material and extraction

The leaves of *A. pubescens* Benth (**Fig 1**) were collected in July 2014 at Covè in Benin and authenticated at the National Herbarium of University of Abomey-Calavi (UAC) where it was kept under voucher AAC 188/HNB.

**Fig 1:**
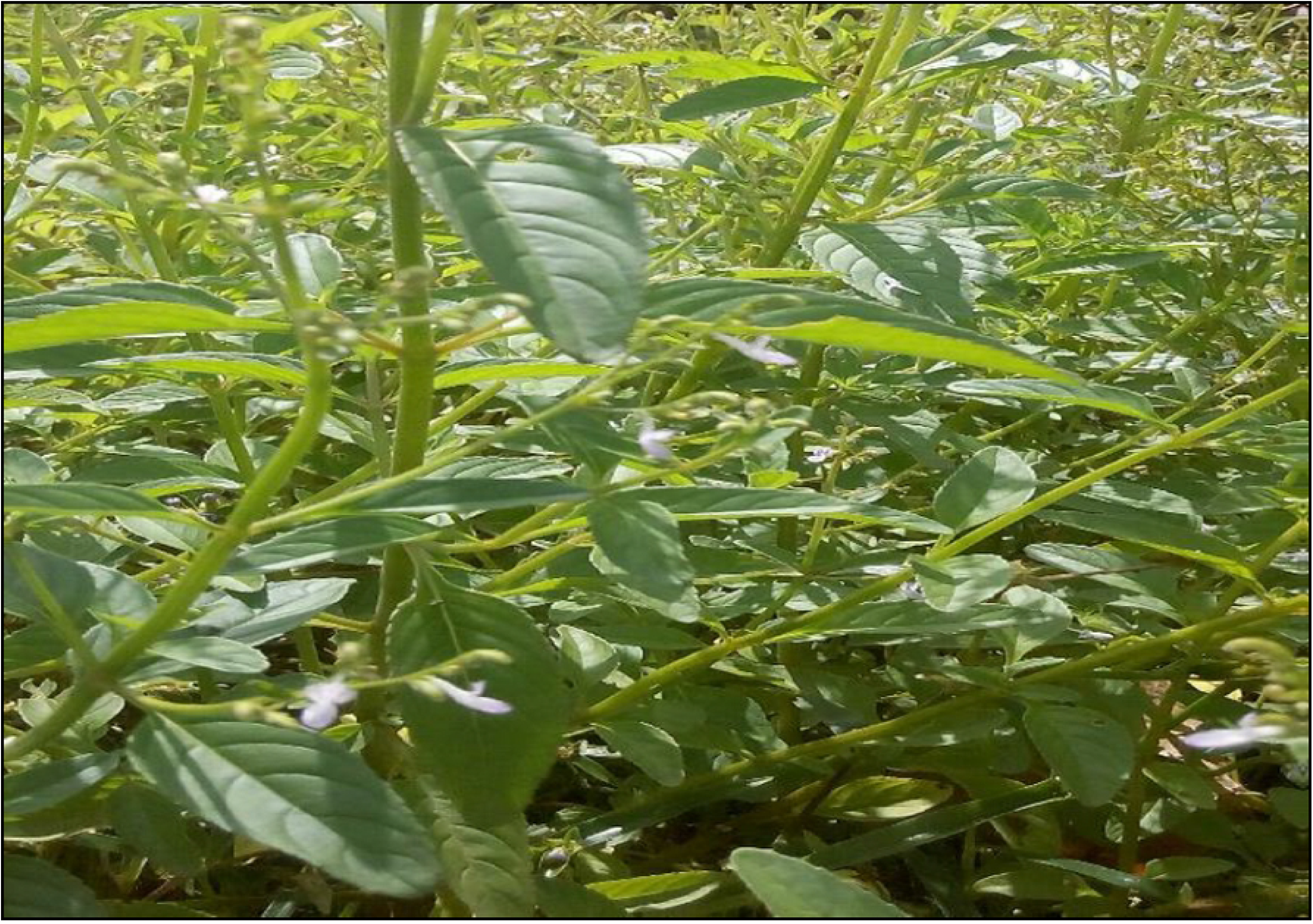
Areal part of *Aeollanthus pubescens* Benth harvested in Benin.

The leaves were dried at 25 °C ±2 °C for 72 hours. Three (03) batches of 200 g of dried leaves were submitted to hydro-distillation in Clevenger apparatus at 100 °C for 2 hours. The distilled oil was dried using anhydrous sodium sulphate and transferred into an airtight amber-coloured vial and stored at 4 °C until further use. The yields were averaged over the three experiments of the plant materials.

### Chemical analysis of *Aeollanthus pubescens* Benth leaves essential oil

#### GC-FID analysis

The essential oil constituents were analysed by capillary gas chromatography coupled with flame ionization detection (GC-FID), equipped with a Supelco SPB-1 column (30 m×0.32 mm i.d, 0.25 µm film thickness). One (1) µL of the essential oil diluted in chloroform were directly injected into the GC. Helium was used as carrier gas with the flow rate of 6 mL/min and the splitting ratio of 1/17. The inlet temperature was 250 °C/min, 200-310 °C at 20 °C/min, and then maintained at 310 °C for 2 min.

#### GC-MS analysis

Capillary gas chromatography coupled with a mass spectrometry detector (GC-MS) was used on a TR-1MS column (30 m x 0.25 mm i.d., 0.25 µm film thickness). An electron impact was used with ionization energy of 70 eV. Helium was used as the carrier gas at a flow rate of 0.6 mL/min, and the splitting ratio 1/17. The temperature settings were as follows: 70-200 °C at 10 °C/min, 200-300 °C at 20 °C/min, and then maintained at 300°C for 1 min. Inlet and MS transfer line temperatures were set at 250 and 320 °C, respectively. All apparatus and accessories were from Thermo Scientific (Courtaboeuf, France) and software controlled data processing (Chromocard and XCalibur). The identification of the essential oil constituents was based on the comparison of their retention times and their Kovats retention indexes relative to (C_8_-C_20_) n-alkanes. Whenever possible, identifications were based on mass spectra of the authentic standard compounds. Otherwise, identifications were performed using published data [26] and comparison with the NIST mass spectral library.

### Mosquito strains

Three *Anopheles gambiae* s.s (*An. gambiae*) laboratory strains that were already available at the insectarium of the laboratory of Vector-Borne Infectious Diseases at the Institut Régional de Santé Publique Alfred Quenum (IRSP-AQ) of the University of Abomey-Calavi in Ouidah (Benin) were used in this study. Kisumu strain originating from Kenya is a reference strain susceptible to all insecticides [27]. Acerkis, which is resistant towards both organophosphate and carbamate based insecticides and is homozygous for (G119S) mutation [28]. Kiskdr, which is homozygous for *kdr*^*R*^ (L1014F) and confers resistance to pyrethroids and DDT [29]. Both AcerKis and Kiskdr were supposed to share the same genetic background as the Kisumu strain but differ by the presence of resistance alleles.

The colonies of each strain were maintained at the insectarium under optimum conditions (25-27°C temperature and 70-80% relative humidity). The third instar larvae, as well as adult females of 3-5 days old of each mosquito strains were used for the bioassays.

### Bioassays

#### Larval bioassay

The larvicidal properties of the essential oil were conducted according to the standard method recommended by the World Health Organization [30] with slight modifications. Since the essential oil does not dissolve in water, six different concentrations (1000, 2000, 3000, 4000, 5000 and 6000 ppm) of the essential oil were prepared in ethanol 96%. Twenty-five (25) third instar larvae of each strain were gently transferred gently into plastic beaker containing 99 mL of water and 1 mL of each prepared concentration was added to obtain test solutions of 10, 20, 30, 40, 50 and 60 ppm respectively. In each round of the experiment, larvae were exposed for 24 hours at 26 ± 2°C without any food provided to them. After exposure, the larval mortality was recorded. Larvae were considered dead when they were not able to move or swim actively when touched. For each strain, four replicates were performed for a total of 100 larvae per concentration. The control group consisted of batches of larvae exposed to water and the solvent alone (ethanol). In total, three different experiments were conducted on three different days.

### Adult Bioassay

#### Impregnation of mosquito nets with the essential oil

Fragments of the non-insecticide treated net (13 cm x 13 cm; 169 cm^2^) corresponding to the surface necessary for the base of the cone were cut and coated with the essential oil. Concentration ranges corresponding to 10, 20 and 30 fold of that of the standard PermaNet 2.0 (55 mg/m^2^ of deltamethrin) were chosen for the impregnation after preliminary doses screening. Then, the essential oil which was yellow with a distinct and sharp odour, was incorporated into the net fragment fibres at 55, 110 and 165 µg/cm^2^.

The masses of essential oil proportional to the net area (169 cm^2^) per concentration were determined: 9.3, 18.6 and 27.9 mg for the impregnation at 55, 110 and 165 ng/cm^2^ respectively. A volume of 1.5 mL of ethanol HPLC grade was poured into a Petri dish containing the mass of essential oil corresponding to a given concentration. After complete dissolution, the fragment of the mosquito net was soaked in the mixture. The impregnated fragment nets were left to dry at room temperature for 5 minutes to allow the essential oil to adhere to the mosquito net and to evaporate the ethanol completely. After drying treated fragment nets were stored in the dark and used 2 to 4 hours later. All coated fragment nets used during the day were treated in the morning at the same time. The net of the same size were also treated with 1.5 mL of solvent and was used as control.

#### Cone test

The cone test was used to assess the effectiveness of the essential oil on the adult mosquitoes. The cone test is an adaptation of the WHO cone bioassay with the following modification: During the assay, the test operator holds a forearm behind the cone to provide a host (**Fig 2**). Unfed 3-5 days old female mosquitoes of Kisumu, Acerkis and Kiskdr strains were used in the test. On the day of testing, female were starved for 4 hours before testing. Groups of five female mosquitoes were placed into plastic cups and moved into the testing room one hour before testing begins to allow the mosquitoes to acclimatise to room conditions. The fragment nets for test or control were placed over a dedicated hole on the Perspex boards and secured using a clear tape. A second Perspex board was laid on the first board creating a test/control net “sandwich” between the two boards. The cone was placed over the net and plugged above with a piece of parafilm. A batch of 5 mosquitoes was transferred into the cone with the operator’s forearm in position. Mosquitoes were then exposed to the fragments for three (03) minutes. Ten (10) replicates of batches of 5 mosquitoes of each strain were run per concentration of impregnated nets.

**Fig 2.**
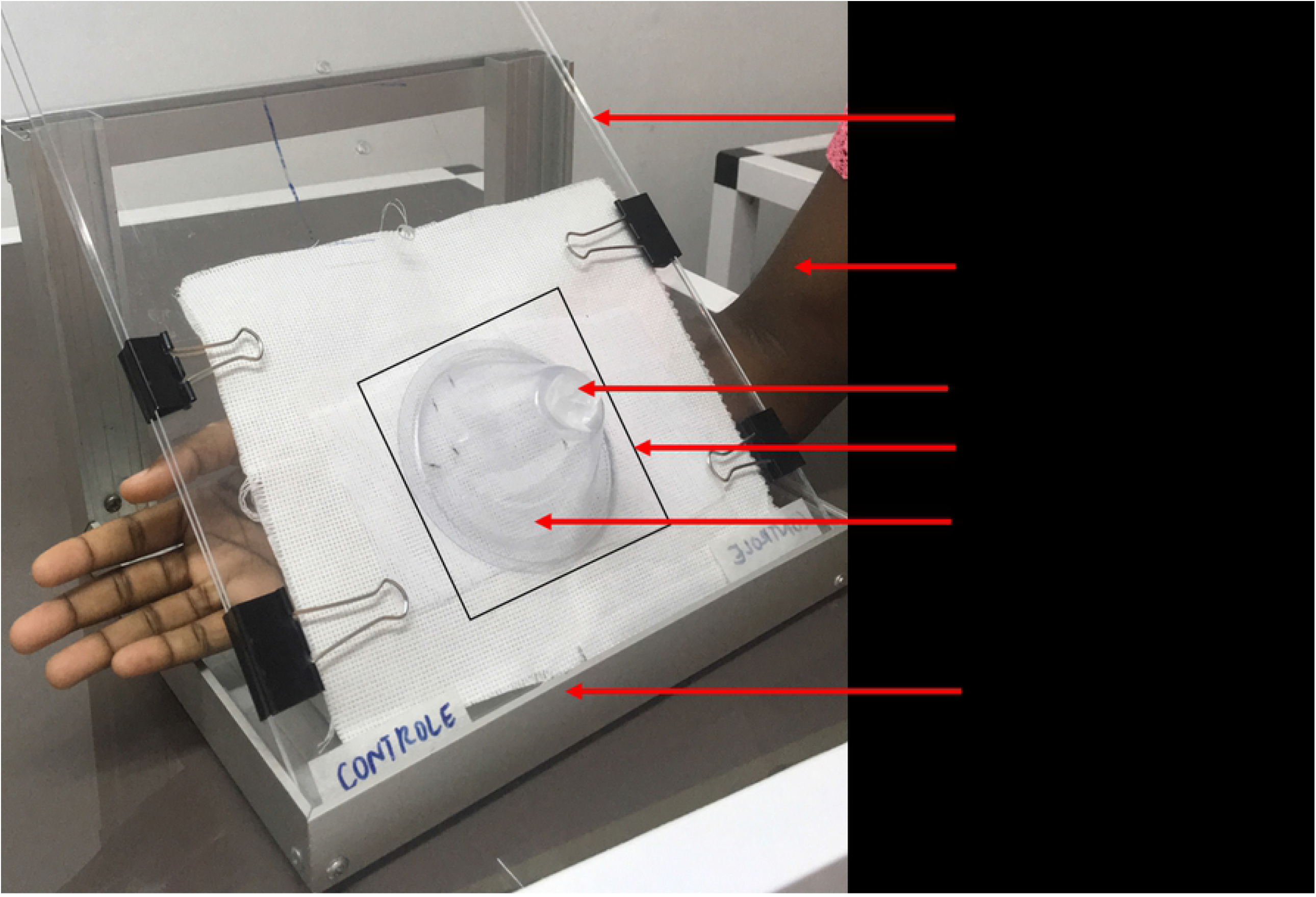
Cone test equipment.

#### Monitoring of the lethal effect of mosquito exposure to the essential oil

After exposure, mosquitoes were removed from the cone, transferred into a recovery cups and provided with 10% of honey solution soaked on a cotton pad. Mosquito knockdown was recorded at 60 minutes post-test. Mosquito mortality was then recorded every day until the death of the last female of each mosquito strain.

### Data analysis

The analysis of dose-mortality responses in larval bioassays was performed using the BioRssay script version 6.2 [31] in R software Version 3.0 [32]. This script computes the lethal concentrations inducing 50% (LC_50_) and 95% (LC_95_) mortality recorded in each strain and the associated confidence intervals; the resistance ratios, i.e. RR_50_ or RR_95_ (LC_50_ or LC_95_ in each strain divided respectively by the LC_50_ or LC_95_ of the reference strain) and their 95% confidence intervals. The times at which 50% or 95% of mosquitoes fell on their back or their side, i.e. knockdown time (KDT_50_ or KDT_95_) and their 95% confidence intervals were estimated after probit regression in R software using the package ‘ecotox’ [33] based on the method described by Finney [34]. The mosquito survival after exposure to the essential oil impregnated net was analysed by Kaplan–Meier survival curves using GraphPad Prism 8.0.2 software (San Diego, California USA). The Log-rank test was performed to evaluate the difference in survival between the strains. All statistical analyses were set at a significance threshold of *p* < 0.05.

## Results

### Extract yield (dry weight %) and Chemical composition of *Aeollanthus pubescens* Benth leaf essential

The percentage yields of essential oil obtained from the hydro-distillation of the leaves of *A. pubescens* was 0.30 ± 0.02 (w/w based on fresh leaves). From the chemical composition of the essential oil of *A. pubescens* Benth (**Table 1**), fourteen compounds were identified, accounting for 98.31% of the crude essential oil’s mass. The essential oil of *A. pubescens* aerial part had higher oxygenated monoterpenes (60.4%) than monoterpene hydrocarbons (22.39%) and sesquiterpene hydrocarbons (15.52%) **(Table 1)**. The major component of the essential oil was carvacrol (51.06%), followed by other components thymol acetate (14.01%), γ-terpinene (10.60%), O-cymene (8.40%) and thymol (5.46%). The percentage of composition of the remaining nine compounds ranged from 0.19 to 2.02% (**Table 1**).

**Table 1:** Chemical composition of the *Aeollanthus pubescens* Benth essential oil.

| Peak No | RI | Components | Area (%) |
| --- | --- | --- | --- |
| 1 | 940 | α-Pinene | 0.58 |
| 2 | 986 | Myrcene | 2.02 |
| 3 | 1005 | Lumicolchicine | 0.19 |
| 4 | 1020 | o-Cymene | 8.40 |
| 5 | 1031 | 1,8-cineole | 0.60 |
| 6 | 1057 | γ-Terpinene | 10.60 |
| 7 | 1088 | Linalool | 0.88 |
| 8 | 1162 | Borneol | 1.40 |
| 9 | 1173 | Terpin-4-ol | 1.60 |
| 10 | 1273 | Thymol | 5.46 |
| 11 | 1284 | Carvacrol | 51.06 |
| 12 | 1359 | Thymol acetate | 14.01 |
| 13 | 1488 | β-Cubebene | 0.24 |
| 14 | 1503 | Acid [(2,4,6-triethylbenzoyl) thio]acetic | 1.26 |
|  |  | Total identified (%) | 98.31 |
|  |  | Sesquiterpenes hydrocarbons | 15.52 |
|  |  | Monoterpenes hydrocarbons | 22.39 |
|  |  | Oxygenated monoterpenes | 60.4 |

RI: relative retention indices as determined on an HP-1 column using the homologous series of n-alkanes.

### Toxicity of the essential oil on *Anopheles gambiae* s.s larvae

We carried out a total of 24 larval bioassays with four replicates for each of the three mosquito strains. The strains Acerkis larvae (LC_50_ = 22.65 ppm) were significantly more susceptible to the essential oil compared to Kiskdr (LC_50_ = 28.37 ppm, *p* < 0.001) and Kisumu (LC_50_ = 29.26 ppm, *p* < 0.001) (**Table 2**). However, Kisumu and Kiskdr larvae susceptibility was not significantly different (*p* = 0.69). All resistance ratio (RRs) calculated were not significantly different to 1 (the RR_50_ was 0.77 and 0.96 in Acerkis and Kiskdr respectively).

**Table 2:** Toxicity against *Anopheles gambiae* larvae after 24 h exposure.

| Mosquito strains | LC<sub>50</sub> (ppm) | 95% C.I | RR<sub>50</sub> | 95% C.I | LC<sub>95</sub> (ppm) | 95% C.I | Slope ±S.E |
| --- | --- | --- | --- | --- | --- | --- | --- |
| Kisumu | 29.26 | 27.73 - 30.73 | - | - | 47.29 | 44.12 - 51.71 | 7.88 ± 0.58 |
| Acerkis | 22.65 | 20.18 - 24.97 | 0.77 | 0.63-0.96 | 44.80 | 39.48 - 53.49 | 5.55 ± 0.57 |
| Kiskdr | 28.37 | 26.84 - 29.83 | 0.96 | 0.79-1.19 | 48.00 | 44.67 - 52.59 | 7.19 ± 0.51 |

LC<sub>50/95</sub>: lethal concentrations; S.E: standard error; C.I: Confidence interval; RR<sub>50</sub> is resistance ratio at LC<sub>50</sub>: LC<sub>50</sub> (resistant strain)/ LC<sub>50</sub> (Kisumu).

### Adulticidal activity of the essential oil

#### Knockdown time of essential oil impregnated net on *Anopheles gambiae* strains

The average time estimated for knockdown 50% (KDT_50_) or 95% (KDT_95_) of adult *An. gambiae* females of each strain decreased with the increasing treatment concentration. The KDT_50_ was less than 4 seconds for all mosquito strains in contact with fragment net treated at 165 µg/cm^2^ (3.77 s for Kisumu; 1.71 s for Acerkis and 2.67 s for Kiskdr), which were significantly lower than that recorded with the lowest essential oil treatment (55 µg/cm^2^) (Kisumu: 22.06 s, *p* < 0.001; Acerkis: 291.72 s, *p* < 0.001; Kiskdr: 591.63s, *p* = 0) (**Table 3**). At the highest treatment concentration (165 µg/cm^2^), both Acerkis and Kiskdr mosquitoes were quickly knocked down (KDT_50_ of 1.71 s, *p* < 0.001 and 2.67 s, *p* < 0.001 respectively) than Kisumu individuals (KDT_50_: 3.77 s).

**Table 3:** Probable time for 50 and 95% knockdown of *Anopheles gambiae s*.*s*. strains per fragment net treatment.

| Strain | Essential oil treatment (µg/cm<sup>2</sup>) | KDT<sub>50</sub> (s) | 95% CI | KDT<sub>95</sub> (s) | 95% CI |
| --- | --- | --- | --- | --- | --- |
| **Kisumu** | 55 | 22.06 | [20.03-23.82] | 45.31 | [40.51-53.26] |
|  | 110 | 4.74 | [4.54 - 4.93] | 6.27 | [5.92 - 6.8] |
|  | 165 | 3.77 | [3.55 - 3.98] | 5.65 | [5.21 - 6.36] |
| **Acerkis** | 55 | 291.72 | [280.64 - 302.38] | 373.70 | [355.49 - 400.70] |
|  | 110 | 4.63 | [4.44 - 4.82] | 6.17 | [5.79 - 6.77] |
|  | 165 | 1.71 | [1.52 - 1.89] | 3.52 | [3.06 - 4.29] |
| **Kiskdr** | 55 | 591.63 | [576.42-607.56] | 813.01 | [777.61-859.02] |
|  | 110 | 197.44 | [185.02-209.26] | 308.68 | [285.79-341.92] |
|  | 165 | 2.67 | [2.46-2.86] | 4.40 | [3.99-5.09] |

(s): second; **KDT<sub>50</sub>** and **KDT<sub>95</sub>**: Knock-down times for 50 and 95% of adult mosquitoes after three minutes of exposure to impregnated fragment net with the essential oil in cone test; **CI**: Confidence Interval.

However, the highest knocked down times values were observed for the Kiskdr (KDT_50_ > 597s) and Acerkis (KDT_50_ >291s) females exposed to the essential oil at 55 µg/cm^2^.

#### Mortality induced by essential oil the impregnated net on *Anopheles gambiae* strains

Overall, the three essential oil treatments had decreased significantly the survival of all mosquito strains after exposure. For the essential oil coating at 165 µg/cm^2^, the longevity of the three mosquito strains decreased significantly from twenty-four (24) days for Kisumu, twenty-five (25) days for Acerkis and twenty-six (26) days for Kiskdr in control groups to respectively one (01) days for Kisumu (*p* < 0.001), two (02) days for Acerkis (*p* < 0.001) and three (03) days for Kiskdr (*p* < 0.001) in exposed groups (**Fig 3C**). With the net treated at 110 µg/cm^2^, Kisumu females longevity was significantly reduced by 21 days compared to that recorded with the 55 µg/cm^2^ treatment (by 14 days; *p* < 0.001) (**Figure 3A and 3B**). With each of these two treatments, no significant effect was observed on the longevity of Kiskdr (*p* = 0.8).

**Fig 3.**
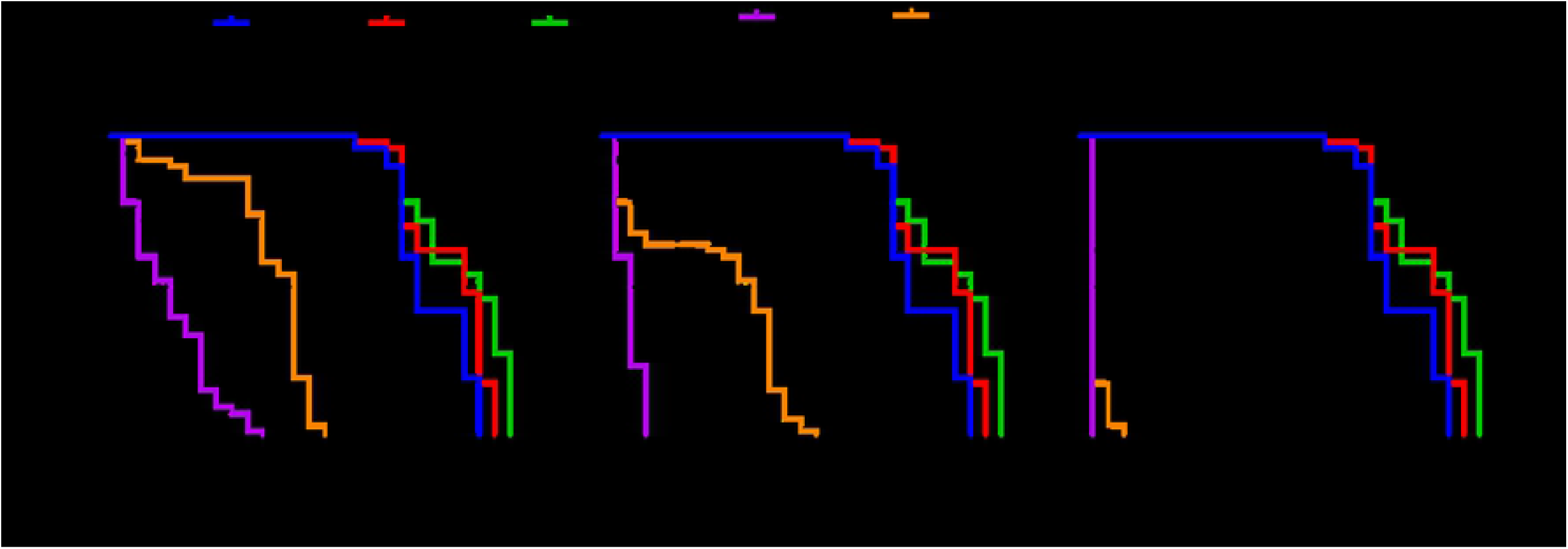
Survivorship of mosquitoes post exposure. Each of mosquito strains was followed up after exposure to fragment net impregnated at 55 µg/cm^2^ (**A**); 110 µg/cm^2^ (**B**) and 165 µg/cm^2^ (**C**).

## Discussion

The increasing number of reports of natural mosquito resistance to the existing synthetic insecticide has strengthened the focus on searching for environmentally-friendly insecticide compounds for vector control strategy. This is a beneficial alternative as essential oils represent a rich source of bioactive compounds that are biodegradable into non-toxic products and due to their natural synergism, they reduce the risk of the development of resistance in the vectors [35]. Besides, essential oils are known as nucleophilic in nature and hinder efficiently with a range of biological processes (metabolic, physiological, biochemical and behavioural) in insects [36–38]. This study is the first report of the larvicidal and adulticidal activity of the *A. pubescens* Benth leaves essential oil on the major African malaria vector *An. gambiae s*.*s*. The insecticidal properties of the essential oil of *A. pubescens* leaves were carried out in the laboratory using immature and adult stages of *An. gambiae* mosquitoes.

The essential oil isolated by hydro-distillation process from the aerial parts of *A. pubescens* collected during this study was found to be a pale yellow liquid, with a yield of 0.30 ± 0.02%. The same oil yield (0.3%) was reported for the fresh aerial material of *A. pubescens* collected from Togo with the same isolation technique [39]. Agbodan et al. [40] have also reported similar oil yield (0.20%) by steam distillation of the fresh leaves of *A. pubescens* collected from Togo. However, a slight increase of the oil yield (1.2 to 1.5%) was reported for the fresh leaves collected in other regions in Benin [25]. These observations suggest that *A. pubescens* oil yield does not depend on the drying step, is neither specific to the extraction technique but may depend on the oil constituents (chemotype). However, the increase in yield of extracted oil could be achieved using the supercritical fluid extraction which will also improve the quality of the extract [41].

The chemical composition analysis of the essential oil of *A. pubescens* aerial parts showed a quantitative variation of all and the major compounds identified. The chemical analysis has displayed the presence of 14 compounds. Carvacrol was the major compound representing 51.06% of the constituents, followed by the thymol acetate (14.01%) and γ-terpinene (10.60%). This oil composition is characteristic of the carvacrol chemotype. Overall, five (05) different chemotypes were identified from the essential oil of *A. pubescens* aerial part from Togo: i) the thymol chemotype: containing 46.3-58% of thymol; ii) the carvacrol chemotype: 58.21% of Carvacrol; iii) the carvacrol and thymol chemotype: 41% of Carvacrol and 27% of thymol, iv) the carvacrol and thymol acetate chemotype: 55.36% of Carvacrol and 35.05% of thymol acetate, and v) the D-fenchone chemotype: 83.69 % of D-fenchone [39,40,42]. However, for the same plant material collected in the central regions of Benin, Alitonou et al. [25] carried out only the thymol (63%) and carvacrol (51.1%) chemotypes. This variability of the chemical composition of essential oils of the same plant material could be due to many factors including: the bioclimate, soil composition, the harvesting period, the geographical location, the degree of maturity of the plant, the seasonal variation and even the plant genetic background [43].

The results from larvae bioassays using the laboratory colonies of *An. gambiae* showed that the essential oil of *A. pubescens* aerial parts is highly active (LC_50_ < 50 ppm) on the specimens of these strains according to the classification of Komalamisra et al. [44]. Surprisingly, the **RR**_**50**_ for Acerkis (0.77) was less than one (01), indicating that Acerkis strain is more susceptible to the oil than Kisumu strain while no apparent difference was found between Kiskdr and Kisumu (**RR**_**50**_ very close to 1). These results suggest that the essential oil had a promising larvicidal property with low LC_50_ values. The significant activity on the resistant strain over the reference susceptible Kisumu mosquitoes indicates that the essential oil does not affect one of the former target sites (Ace-1^R^ and Kdr^R^ (L1014F) alleles) represented in the corresponding strain. Therefore, it could be implied that its mode of action is different from that of pyrethroids, organophosphates and carbamates. However, in the context of the increasing insecticide resistance in natural mosquito populations, it will be interesting to investigate the bioactivity of this essential oil on field-collected larvae and field caught-adults.

Other biological activities of *A. pubescens* essential oil were reported including antioxidant [25,40], antibacterial [45], but no report was made on its mosquitocidal activity so far. In addition, there is still a lack of information on the bioinsecticidal property of the other plant species belonging to the genus *Aeollanthus*. However, previous studies have investigated the larvicidal activity of the essential oil of plants belonging to the same family (Lamiaceae). Indeed, Tchoumbougnang et al. [46] showed that *Ocimun canum, Ocimum gratissimum* and *Thymus vulgaris* displayed respectively LC_50_ value of 201, 180 and 119 ppm on field-collected *An. gambiae* larvae. These values are higher than those recorded in our study. This variation could be due to the difference in the oils chemical composition and the genetic background of the larvae strains used. Moreover, essential oils from other *Lamiaceae* species (*Plectranthus amboinicus* and *Plectranthus mollis*) were found to be active against *Anopheles stephensi* larvae with LC_50_ value less than 50 ppm [47,48]. These findings suggest that essential oils from *Lamiaceae* plant species could be a potential source of environmental eco-friendly mosquitocidal agents. In our study, *A. pubescens* essential oil was dominated by monoterpenes (oxygenated and hydrocarbons) which represented 82.79% of the oil. Other plants species with similar major constituents have been reported to be active against *An. gambiae* larvae. Ollengo et al. [49] reported that *Clausena anisata* containing 56.7% of monoterpenes revealed a potential larvicidal activity against *An. gambiae* (LC_50_ = 75.96 ppm) [50]. Also, Wangrawa et al. [51] demonstrated that *Lantana camara* essential oil with 70.5% of monoterpenes showed differential larval mortalities on both the laboratory and the field strains of *An. gambiae*. The high proportion of monoterpenes in the essential oil could be correlated to the observed bioactivity. Indeed, many studies reported the larvicidal effect of monoterpenes against mosquitoes strains [52–54]. Therefore, it would be interesting to evaluate further the toxicity of the monoterpenes isolated from *A. pubescens* areal part on mosquito larvae in both laboratory and field trials.

We assume that Carvacrol, the main compound (51.06%) found within the monoterpenes in our *A. pubescens* oil extract, might be the main contributor of the observed activity against *An. gambiae* larvae. Carvacrol is well known for its larvicidal property against both *Anophelinae* (*Anopheles stephensi, Anopheles subpictus*) [55] and *Culicidae* (*Aedes aegypti, Culex quinquefasciatus* and *Culex tritaeniorhynchu*) [55–58].

Essential oils are mixtures of volatile compounds and due to the antagonistic or synergistic phenomena, the bioactivity of the crude oil extract in some cases is lower or higher than those of purified compounds. For instance, Evergetis et al. [59] demonstrated that larvicidal activity of the essential oil of *Origanum vulgare* ssp against *Aedes albopictus* (LC_50_ = 30.1ppm) is lower than that of its major component, the pure Carvacrol (LC_50_ = 13.1 ppm), accounted for 88.7% of the oil. The same trend was noticed with the leaf essential oil of *Coleus aromaticus* which displayed lower toxicity than its major component Carvacrol against *Anopheles stephensi* larvae [60]. This open perspectives for further investigations to evaluate the larvicidal efficacy of the Carvacrol in comparison to that of the crude oil extract. It is well known that monoterpenes from essential oils could act by absorption through the cuticle, or via the respiratory tract or by ingestion via the gastrointestinal tract [61–64]. Besides, several monoterpenes were reported to target primarily the cholinergic, octopamenergic and GABA neurosystems in insects [65]. One or a combination of these mechanisms might be the pathway of mortality induction by the *A. pubscens* oil. In this study, the reference resistant strain Acerkis larvae haboring *ace-1*^*R*^ allele coding for the insensitive acetylcholinesterase enzyme was the most susceptible to our essential oil. This indicates that the essential oil overcomes the target site modification resistance mechanism and therefore appears as a hopeful alternative tool for vector control programs.

Among the vector life-history traits, mosquito survival is strongly associated with the malaria transmission intensity [66]. Thus, this study also investigated the effect of exposure to various doses of *A. pubescens* oil on adults *An. gambiae* survival. The *A. pubescens* essential oil reduced significantly the lifespan of the three mosquitoes strains exposed to the fragment net at 165 µg/cm^2^. None of the three mosquito strains was able to survive after 72 hours. This observation suggests that even the resistance mosquitoes (Acerkis and Kiskdr) could not survive long enough to allow the extrinsic incubation period of the *Plasmodium* parasites if they ingested a gametocyte infected blood meal. Overall, the drastic reduction in daily survival of mosquitoes observed with the oil treatment at 165 µg/cm^2^ would contribute to a reduction in vectorial capacity in a typical endemic setting and therefore will lead to a reduction in parasite transmission according to the Ross-MacDonald model [67]. This is a promising finding for the management of the resistant malaria-transmitting vectors. Spray-type solution formulations could be made towards the development of botanical insecticides for the use of an integrated approach with the existing conventional vector control strategies. However, the results were obtained using mosquito strains in which only one resistance mechanism is present. It will be interesting to evaluate further the survivorship of the natural mosquito populations where several resistance mechanisms could coexist. The susceptibility of *Plasmodium* infection following essential oil exposure is also another promising parameter to be evaluated.

During the experiment, we observed that the legs of mosquitoes were detached from their bodies when exposed to the net coated at 165 µg/cm^2^. To our knowledge, this phenomenon has not been observed yet. This effect of the essential oil may be caused by its main constituent, the Carvacrol. However, such mechanism is so far unexplored. Possible neurotoxicity in insects could have been easily overlooked. Thus, investigations around the mechanisms used by this compound is urgently sought. The mosquito legs loss suggests that the essential oil could interfere with the insect locomotor system and even the nervous system leading to the death in the following days.

At the doses 55 and 110 µg/cm^2^ of the essential oil, the lifespan of mosquitoes is fourteen (14) days maximum for both resistant Acerkis and Kiskdr strains. During this time period, mosquitoes might still be able to reproduce. Therefore, further studies are needed to assess the blood-feeding success, the fecundity and the fertility of mosquitoes following exposure. This could lead to highlight putative detrimental effects of the essential oil exposure that could also hamper the vectorial competence of the mosquitoes.

## Conclusion

The present study paves the way to develop new and safer natural insecticide and repellent against malaria mosquito vectors. The *A. pubescens* Benth essential oil is proved to be an efficient larvicide and adulticide against malaria vector *An. gambiae*. This opens the perspectives for implementing sustainable control of mosquito populations that are resistant to the current existing synthetic insecticides. The larval and adult vector control with the essential oil could be considered in an integrated fashion to the existing malaria control strategies. Further studies are needed to help in designing *A. pubescens* essential oil formulation that would potentially increase its efficacy on *An. gambiae* and its cost-effectiveness.

## Acknowledgments

The authors thank Geraldine Foster from Liverpool School of Tropical Medicine (LSTM) for the material support. We are grateful to Ms Marie Joelle Fanou for her assistance in insectary work; Mrs Laurette Djossou for her help during data acquisition; Mrs Emilienne Fiogbe and Mr Jean-Louis Amoussou for their helpful proofreading of the manuscript. The authors also acknowledge Wellcome Trust for financial support to LSD (grant 109917/Z/15/Z).

## Author Contributions

### Conceived and designed the experiments

Roméo Barnabé Bohounton, Luc Salako Djogbénou, Pierre Villeneuve, Fidèle Paul Tchobo.

### Performed the experiments

Roméo Barnabé Bohounton, Pierre Marie Sovegnon, Bruno Barea.

### Data analysis

Roméo Barnabé Bohounton, Oswald Yédjinnavênan Djihinto, Oronce Sedjro-Ludolphe Dedome.

### Writing – original draft

Roméo Barnabé Bohounton, Oswald Yédjinnavênan Djihinto.

### Writing – review & editing

Roméo Barnabé Bohounton, Oswald Yédjinnavênan Djihinto, Aristide Adomou, Luc Salako Djogbénou, Fidèle Paul Tchobo.

